# Human placental trophoblasts support sustained *Treponema pallidum* replication and reveal new candidate host pathways implicated in congenital syphilis

**DOI:** 10.64898/2026.05.10.724106

**Authors:** Eliza R McColl, Diane G Edmondson, Bridget D DeLay, Karan G Kaval, Steven J Bark, Steven J Norris, Indira U Mysorekar

## Abstract

Congenital syphilis is a leading cause of preventable stillbirth, yet the mechanisms that enable *Treponema pallidum* subsp. *pallidum,* the etiological agent of syphilis, to traverse and persist within the human placenta are virtually unknown. This knowledge gap reflects, in part, the fastidious nature of *T. pallidum*, which has historically only been propagated in rabbit epithelial cells. Here, we demonstrate that *T. pallidum* replicates and can be propagated long-term in human placental trophoblast models. We identified three human trophoblast cell lines, JEG-3, BeWo, and HTR-8, that sustained robust *T. pallidum* replication for over 55 days. Confocal imaging with GFP-expressing *T. pallidum* demonstrated bacterial adherence to trophoblast cultures. To identify human placental pathways that may support *T. pallidum* replication and persistence, we performed bulk transcriptomic profiling of the three trophoblast lines co-cultured with *T. pallidum* for 3 or 7 days. Our findings revealed conserved host responses involving increased cholesterol synthesis, suppressed type I interferon signaling, and disrupted extracellular matrix organization. These data implicate host metabolic rewiring, innate immune attenuation, and extracellular matrix remodeling as candidate pathways that may promote *T. pallidum* invasion, replication and persistence within the placenta. Together, these models represent the first human placental systems capable of supporting efficient, long-term *T. pallidum* growth and provide a foundation for mechanistic studies of congenital syphilis pathogenesis.

## Introduction

Congenital syphilis, caused by vertical transmission of the bacterial spirochete *Treponema pallidum*, is the second leading cause of preventable stillbirth worldwide.^1^ Surviving infants experience symptoms such as osteochondritis or periostitis, neurosyphilis, rashes, hepatobiliary dysfunction, anemia, and hearing loss.^2^ While rates of vertical transmission of *T. pallidum* reach up to 100% in the third trimester,^2^ prenatal treatment with benzathine penicillin G is highly effective at preventing congenital syphilis (98.2%).^3^ However, inadequate access to prenatal care and screening, diagnostic challenges, and intermittent penicillin shortages have prevented global elimination of congenital syphilis.^4^ Indeed, cases have skyrocketed over 10-fold in the past decade^5^ and over 700,000 cases are estimated to occur annually worldwide.^1^ These challenges underscore the need to define biomarkers of placental infection and actionable host-pathogen pathways that could improve detection and prevention of vertical transmission.

A major barrier to developing such interventions is the limited mechanistic understanding of how *T. pallidum* impacts the human placenta during vertical transmission. Although *T. pallidum* is known to infect and cross the placenta, causing histopathological changes including placentomegaly, villous enlargement, necrotizing funisitis, and acute or chronic villitis,^6–9^ the cellular and molecular events that permit placental invasion, persistence, and fetal dissemination remain unknown. Once in the fetal circulation, *T. pallidum* disseminates systemically and causes widespread fetal injury. However, how this pathogen breaches the placental barrier, establishes infection within placental tissues, and triggers placental and fetal pathology remains poorly understood. This knowledge gap reflects both the lack of animal models that mimic congenital syphilis during human pregnancy and the technical challenges associated with long-term *T. pallidum* culture.^10^

The establishment of long-term continuous culture conditions for *T. pallidum* in 2018 was a major advance for syphilis research.^11^ This breakthrough enabled sustained *in vitro* propagation of this fastidious pathogen in the cottontail rabbit epithelial cell line Sf1Ep in specialized *T. pallidum* Culture Medium 2 (TpCM2) at 34°C under microaerobic conditions. However, because this system relies on rabbit epithelial cells, its ability to model human tissue-specific host-pathogen interactions remains limited. Although *T. pallidum* replication in a human foreskin fibroblast cell line was reported in March 2026,^12^ sustained replication in human placental cells has not been established. This gap has severely limited mechanistic studies of congenital syphilis and constrained efforts to identify placental biomarkers and host pathways that could be targeted to prevent vertical transmission. We therefore sought to determine whether human placental cell lines could support long-term *T. pallidum* growth *in vitro*, which would provide tractable models to define host-pathogen interactions at the maternal-fetal interface and study mechanisms of congenital syphilis.

Originally isolated from choriocarcinomas, JEG-3^13^ and BeWo^14^ are widely used models of cytotrophoblasts (CTBs), trophectoderm-derived cells which fuse to form the maternal-facing continuous barrier of the placenta, the syncytiotrophoblast (STB).^15^ HTR-8/SVneo cells (referred to here as HTR-8)^16^ are typically used to model extravillous trophoblasts (EVTs) which invade the maternal decidua to anchor the placenta.^15^ Here, we report that all three of these human placental cell lines support continuous, sustained *T. pallidum* replication at rates comparable to the Sf1Ep model, thus establishing the first *in vitro* models of *T. pallidum* in the placenta. Transcriptional profiling of *T. pallidum*-infected trophoblasts revealed conserved changes in genes associated with cholesterol metabolism, extracellular matrix organization, and suppression of interferon signaling, providing several new avenues for research into potential mechanisms used by *T. pallidum* to invade and persist in the placenta. As the first human trophoblast cell lines found to support *T. pallidum* replication, these models serve as unprecedented tools to study mechanisms underlying *T. pallidum* pathogenesis during congenital syphilis.

## Results

### Human placental cells support sustained growth and surface adherence of T. pallidum

We first asked whether human placental cell lines could support *T. pallidum* replication under conditions used for the standard Sf1Ep rabbit epithelial cell line. JEG-3, BeWo, and HTR-8 cell lines cultured in TpCM2^11^ were all able to support continuous replication of the *T. pallidum* Nichols strain during the 55-day observation period (**Fig. 1A-C**). While all three cell lines supported *T. pallidum* growth at rates similar to Sf1Ep cells, average generation times in JEG-3 were slightly longer (44 ± 4 h) than in Sf1Ep (37 ± 3 h) (**Fig. 1D**). In contrast, generation times in HTR-8 (36 ± 2 h) and BeWo (40 ± 3 h) were closer to that in Sf1Ep (**Fig. 1E,F**). Together, these data establish JEG-3, BeWo and HTR-8 cells as human placental models that sustain long-term *T. pallidum* replication and enable investigation of host-pathogen interactions relevant to congenital syphilis.

**Figure 1:**
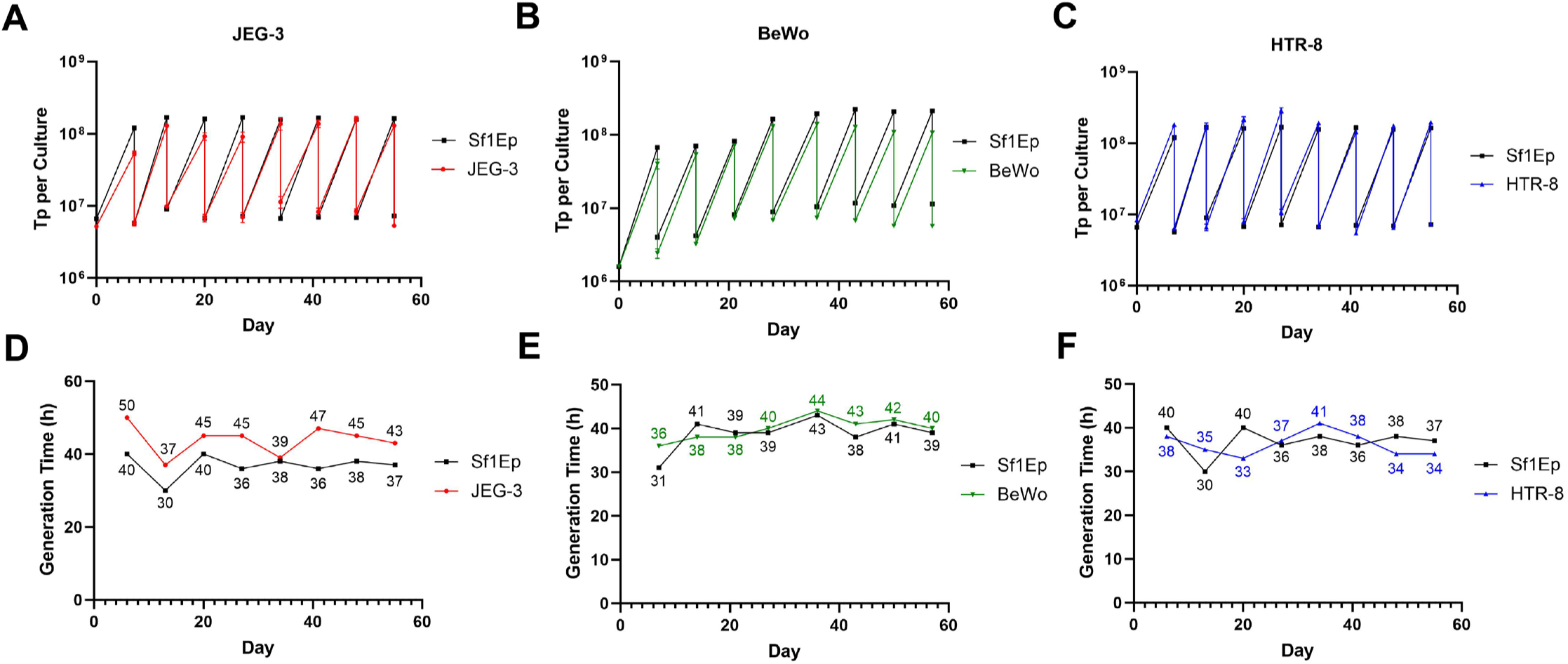
Trophoblast cell lines facilitate efficient *in vitro T. pallidum* replication. Data for JEG-3 (red), BeWo (green), and HTR-8 (blue) are shown relative to standard Sf1Ep cultures (black). (A-C) Sawtooth graphs documenting *T. pallidum* counts per culture over the course of 55 days, with subculture occurring every 6-7 days. (D-F) Average generation time for doubling of *T. pallidum* counts as calculated at time of subculture. n=3 biological replicates, represented as mean ± standard error.

Next, to visualize interactions between placental cell lines and *T. pallidum*, JEG-3, BeWo, and HTR-8 cells were inoculated with a GFP-tagged *T. pallidum* Nichols strain.^22^ Similar to Sf1Ep cells, GFP-labelled *T. pallidum* associated with three placental cell lines (**Fig. 2A**). Three-dimensional imaging suggested that spirochetes localized primarily along the external surface of placental cells and Sf1Ep cells, consistent with surface adherence (**Fig. 2B, Supp. Videos 1-4**).

**Figure 2:**
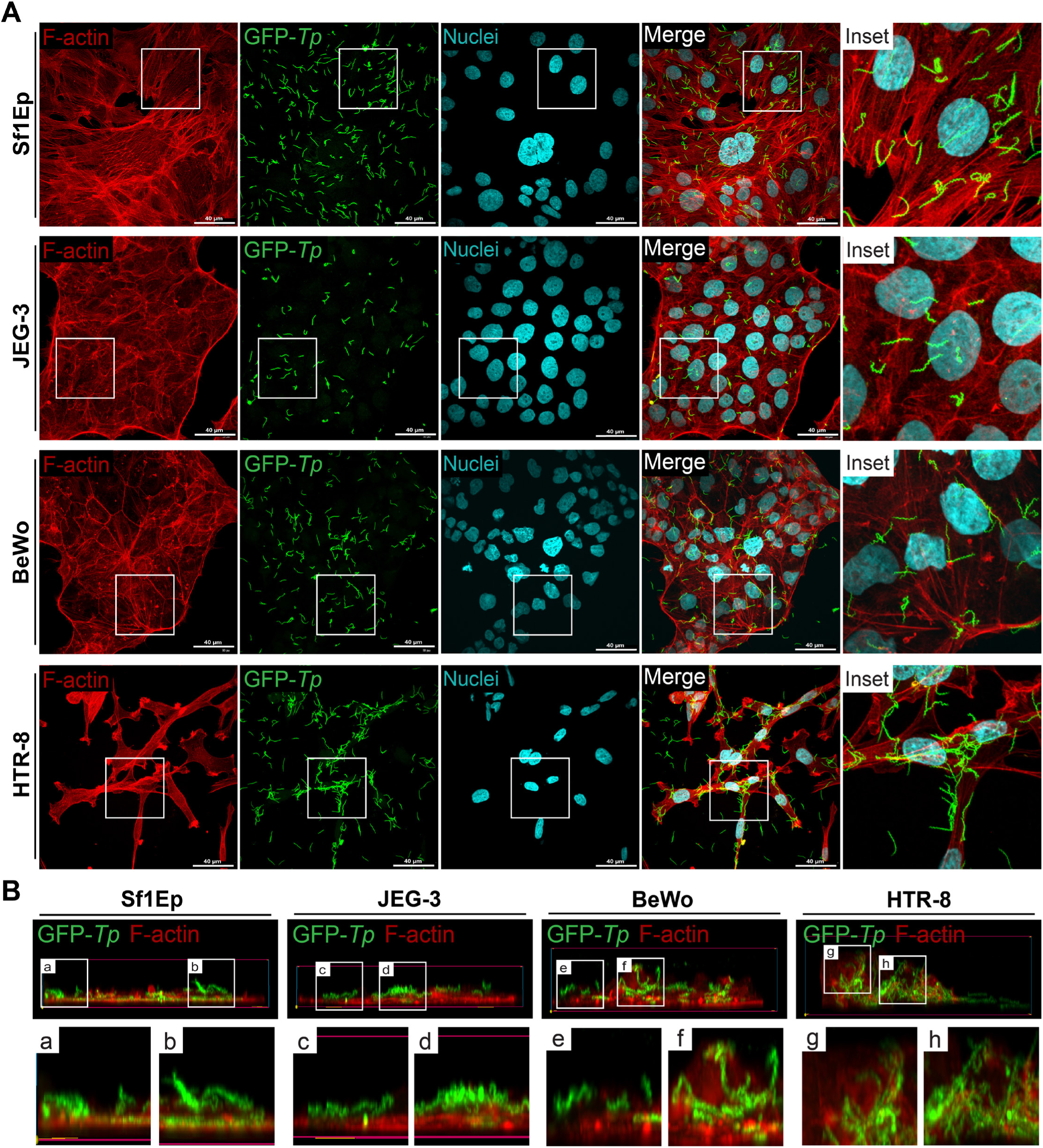
GFP-tagged *T. pallidum* adheres to Sf1Ep and trophoblast cell lines. Cell lines were co-cultured with GFP-tagged *T. pallidum* (GFP-*Tp*, green) for seven days prior to immunofluorescence staining and confocal imaging. Cell margins are stained with F-actin (red) and nuclei are stained blue. (A) GFP-*Tp* adheres to the surface of all four cell lines examined. Scale bar = 40 µm. (B) Lateral views show that GFP-*Tp* appears to colonize the outer surface of cells.

### T. pallidum co-culture remodels host transcriptional programs in human placental cells

To begin defining the host pathways engaged during *T. pallidum* infection, we conducted bulk RNA sequencing on all three cell lines co-cultured with *T. pallidum* for either 3 or 7 days. Co-culture with *T. pallidum* induced robust changes in gene expression of all three cell lines, with distinct differences in transcription between control and *T. pallidum*-infected samples (**Fig. 3**). After three days of co-culture, a total of 281 (233 up, 48 down), 1849 (1260 up, 589 down), and 1767 (1180 up, 587 down) genes were significantly dysregulated (Log_2_FoldChange ≥ |0.5|) in JEG-3, BeWo, and HTR-8 cells, respectively. More extensive changes were observed after 7 days of co-culture, with 1337 (1094 up, 243 down), 1582 (1051 up, 531 down), and 4998 (2706 up, 2292 down) significantly dysregulated genes in JEG-3, BeWo, and HTR-8 cells. For a full list of significantly dysregulated genes, see **Supp. File 2**. These results demonstrate that *T. pallidum* infection significantly disrupts the transcriptome of human placental cells.

**Figure 3:**
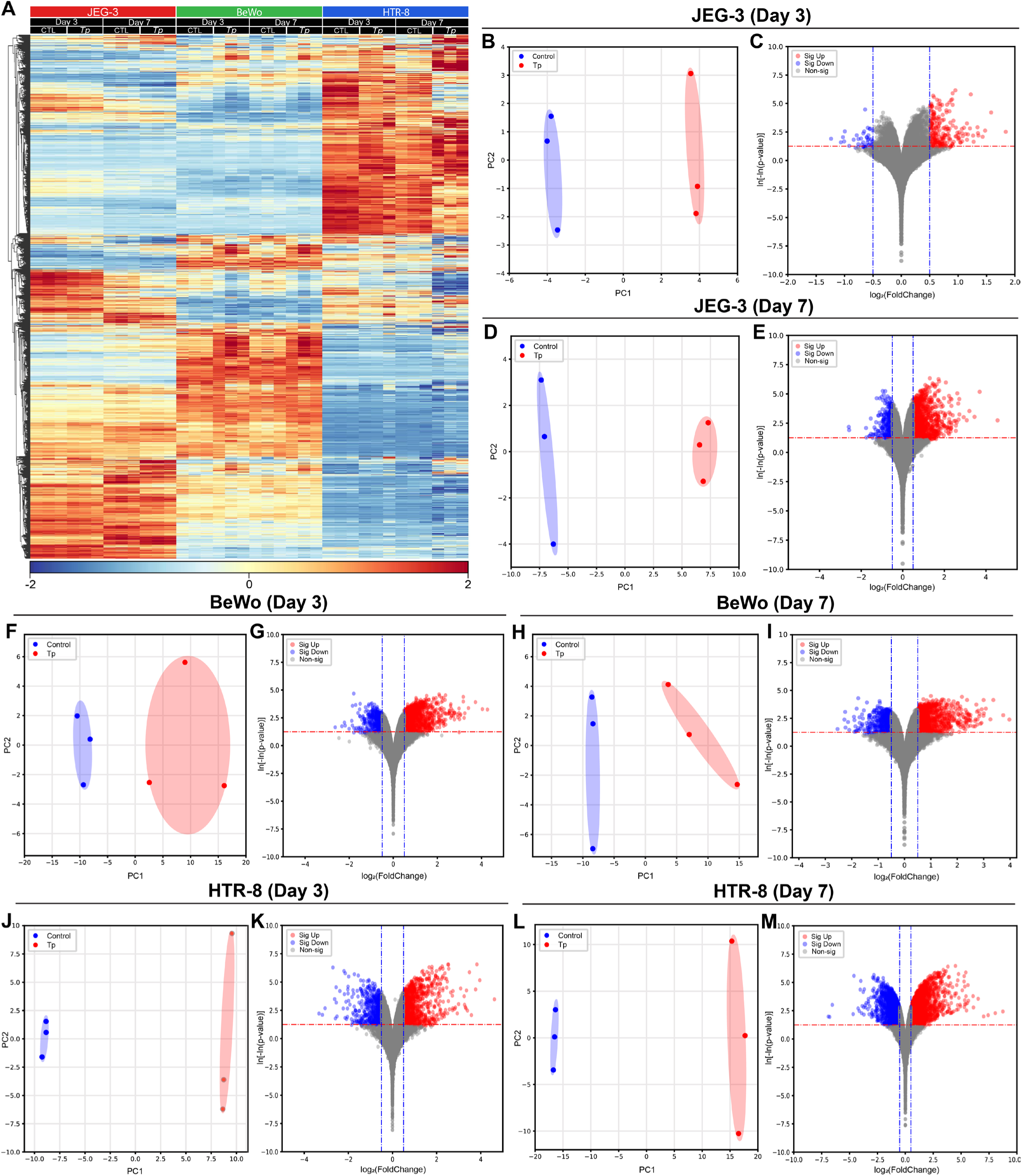
Co-culture with *T. pallidum* elicits widespread changes in gene expression in trophoblast cell lines. JEG-3, BeWo, and HTR-8 cells co-cultured with *T. pallidum* for 3 or 7 days were subjected to bulk RNAseq. (A) Heatmap showing relative changes in gene expression across all three cell lines at both timepoints. Scale refers to z-score. (B,D,F,H,J,L) PCA plots showing separation of transcriptional profiles of control (blue) and *T. pallidum*-infected (red) trophoblasts. (C,E,G,I,K,M) Volcano plots (log_2_FoldChange vs ln[-ln(p-value)]) showing significantly dysregulated transcripts in *T. pallidum*-infected cells relative to controls. Significantly upregulated genes are shown in red, significantly downregulated genes are in blue, and non-significant genes are in grey. Significance was defined as p<0.05 (cutoff indicated by red dotted lines) and log_2_(FoldChange) ± 0.5 (cutoff indicated by blue dotted lines).

### Conserved host transcriptional responses identify placental pathways disrupted by T. pallidum

To identify consistent and conserved transcriptional programs that may be co-opted or disrupted by *T. pallidum* in placental trophoblasts, we searched for transcripts that were significantly altered across all three cell lines and performed pathway enrichment analysis for Gene Ontology biological processes. After 3 days of co-culture, the three cell lines shared 52 significantly upregulated genes (**Fig. 4A, Supp. File 3**). Pathway enrichment analysis of this subset of genes revealed significant enrichment of processes related to cholesterol metabolism, namely secondary alcohol biosynthetic process, cholesterol biosynthetic process, sterol biosynthetic process, cholesterol metabolic process, and cholesterol import (**Fig. 4B, Supp. File 4**). Only one gene, sideroflexin 2 (*SFXN2*), was commonly downregulated in all three cell lines on day 3 (**Fig. 4C**).

**Figure 4:**
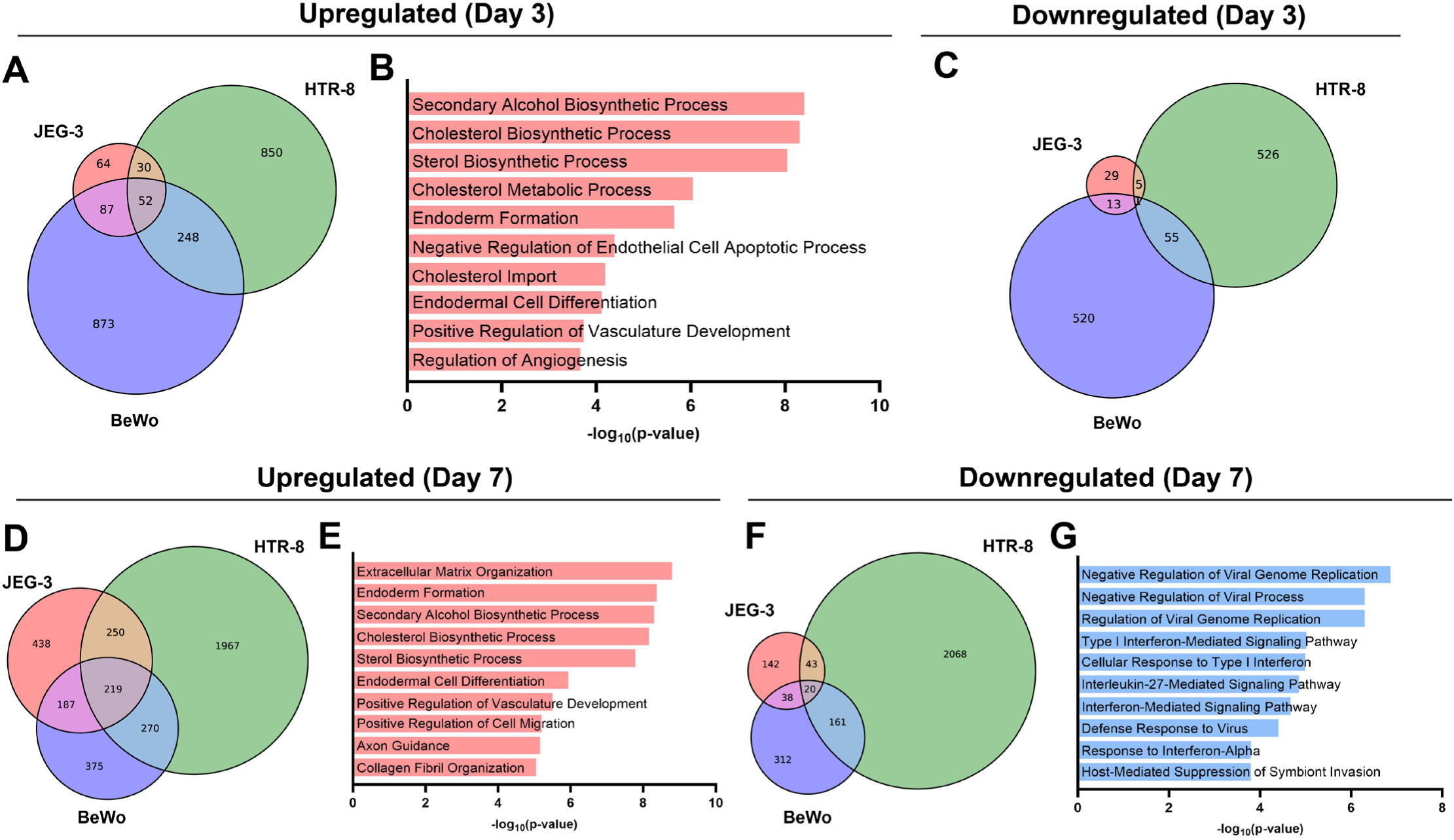
Pathway enrichment analysis of genes similarly dysregulated genes across trophoblast cell lines reveals biological processes disrupted by *T. pallidum*. (A,C,D,F) Venn diagrams showing degree of overlap of significantly up- or downregulated genes between JEG-3 (red), BeWo (blue) and HTR-8 (green) cells co-cultured with *T. pallidum* for 3 or 7 days. (B,E,G) Top 10 most significantly enriched GO biological pathways derived from genes commonly up-(red) or downregulated (blue) across the three trophoblast cell lines.

By day 7, the three placental cell lines shared 219 significantly upregulated genes (**Fig. 4D, Supp. File 3**). Consistent with the day 3 response, these genes remained enriched for cholesterol metabolism, with additional enrichment of pathways related to extracellular matrix organization and cell movement (extracellular matrix formation, positive regulation of cell migration, and collagen fibril organization) (**Fig. 4E, Supp. File 4**). Finally, there were 20 significantly downregulated genes common to all three cell lines on day 7 (**Fig. 4F, Supp. File 3**). Interestingly, these genes corresponded overwhelmingly to immune and pathogen invasion-related pathways, including negative regulation of viral processes and genome replication, type I interferon (IFN)-mediated signaling, interleukin (IL)-27-mediated signaling, defense response to virus, and host-mediated suppression of symbiont invasion. (**Fig. 4G, Supp. File 4**). Together, these analyses identify conserved transcriptional responses in *T. pallidum*-exposed placental cells, implicating cholesterol metabolism, extracellular matrix remodeling and attenuation of interferon-associated host defense programs as candidate pathways in placental colonization and persistence.

### T. pallidum dysregulates trophoblast expression of genes involved in cholesterol metabolism, interferon signaling, and extracellular matrix organization

To confirm alteration of the conserved host pathways nominated by RNA-seq, we examined representative genes involved in cholesterol metabolism, interferon signaling and extracellular matrix remodeling across *T. pallidum*-exposed trophoblast cultures. Given that day 7 had the highest degree of overlap of dysregulated genes between cell lines, qRT-PCR was performed on day 7 samples (**Fig. 5**). In addition to comparisons between control and live *T. pallidum*-infected cells, we determined whether trophoblast transcriptional remodeling required active spirochete metabolism and replication, rather than exposure to bacterial material alone, by profiling cells co-cultured with heat-killed *T. pallidum*.

**Figure 5:**
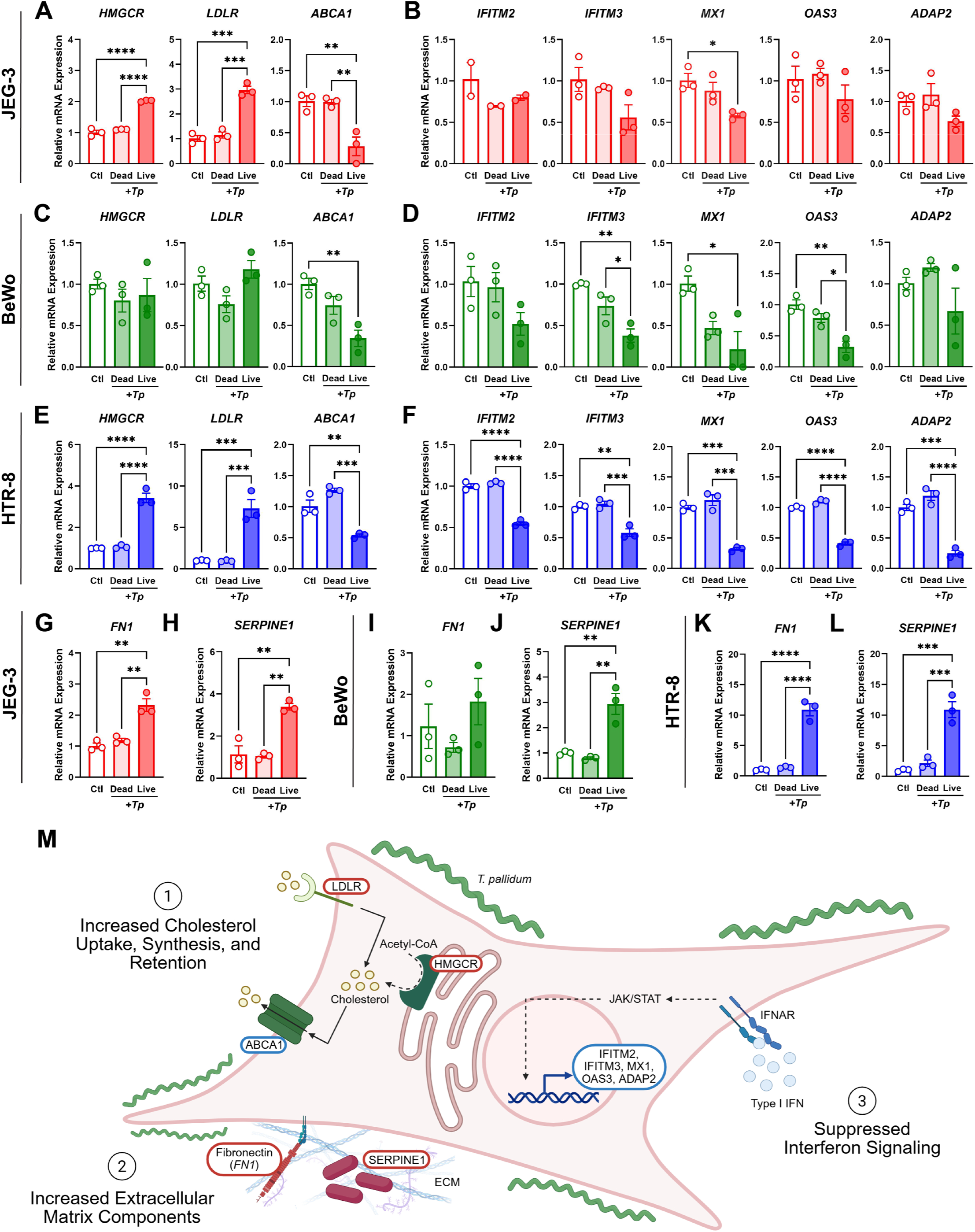
Validation of transcriptional changes in trophoblast cell lines co-cultured with *T. pallidum* for 7 days. Genes that were significantly up- or downregulated across all three cell lines in RNAseq results were subjected to validation with qRT-PCR. Graphs depict relative mRNA expression of genes involved in (A,C,E) cholesterol metabolism, (B,D,F) interferon signaling, and (G-L) extracellular matrix organization in JEG-3 (red), BeWo (green), and HTR-8 (blue) cells. Blank bars correspond to uninfected controls (Ctl), lightly shaded bars correspond to co-culture with heat-inactivated *T. pallidum* (dead), and darker bars correspond to co-culture with live, replication-competent *T. pallidum* (live). Data is shown as mean ± standard error of the mean, n=3 biological replicates. (M) Schematic of a trophoblast cell infected with *T. pallidum*, depicting three proposed disrupted pathways based on transcriptomic analysis. Genes of interest are outlined in red if significantly increased by *T. pallidum*, and blue if significantly decreased. ECM = extracellular matrix, IFNAR = interferon α/β receptor, JAK/STAT = Janus kinase/signal transducer and activator of transcription pathway.

We first assessed transcript levels of three of the genes associated with cholesterol metabolism that were significantly altered across all three cell lines in RNAseq. qRT-PCR confirmed significantly increased mRNA levels of 3-hydroxy-3-methylglutaryl coenzyme reductase (*HMGCR*), a rate-limiting enzyme in cholesterol biosynthesis, and low-density lipoprotein receptor (*LDLR*), which facilitates cellular uptake of cholesterol, in both JEG-3 and HTR-8 cells (**Fig. 5A,C,E**). In contrast, transcript levels of *ABCA1*, a cholesterol efflux transporter, were significantly reduced across all three cell lines. Together, this suggests a conserved mechanism of promoting increased biosynthesis, uptake, and retention of intracellular cholesterol in *T. pallidum*-infected trophoblasts. The absence of differences between controls and those cultured with heat-inactivated *T. pallidum* further suggests that these changes are dependent on active growth and replication of *T. pallidum*, rather than just the presence of spirochete pathogen-associated molecular patterns (PAMPs).

Given the downregulation of IFN-related signaling in the pathway enrichment analysis, we next measured transcript levels of several IFN-stimulated genes (ISGs) that were significantly decreased in RNAseq results. mRNA expression of MX dynamin-like GTPase 1 (*MX1*) was significantly decreased across all three cell lines (**Fig. 5B,D,F**). Interferon-induced transmembrane protein 3 (*IFITM3*) and 2’-5’-oligoadenylate synthetase 3 (*OAS3*) were significantly downregulated in both BeWo and HTR-8, with HTR-8 exhibiting additional significant downregulation of interferon-induced transmembrane protein 2 (*IFITM2*) and ArfGAP with dual pH domains 2 (*ADAP2*).

While some genes were slightly decreased by co-culture with heat-inactivated *T. pallidum* (i.e. *IFITM2* in JEG-3 and *IFITM3* and *MX1* in BeWo), differences between controls and heat-inactivated *T. pallidum* groups were not statistically significant for any genes examined. Together, these data suggest that ISG suppression depends on live *T. pallidum* replication or active host-pathogen interactions, rather than exposure to bacterial PAMPs alone.

RNA-seq also identified two additional genes of interest that were significantly dysregulated across all three placental cell lines. The first, fibronectin 1 (*FN1*), is a known binding target of *T. pallidum*^23^ and was significantly increased (as measured by qRT-PCR) in JEG-3 and HTR-8, with a similar trend observed in BeWo cells (**Fig. 5G,I,K**). The second was *SERPINE1*, which encodes plasminogen activator inhibitor-1 (PAI-1), and was previously reported to be significantly increased in human cerebral brain microvascular endothelial cells exposed to *T. pallidum* for 24 hours.^24^ Consistent with RNAseq results, which identified significant upregulation of *SERPINE1* mRNA across all three cell lines, qRT-PCR found a significant and robust increase in *SERPINE1* in live *T. pallidum*-treated JEG-3, BeWo and HTR-8 cells (**Fig. 5H,J,L**). Together, these results validate conserved transcriptional responses among trophoblast cells in response to *T. pallidum* infection and replication (**Fig. 5M**).

*SERPINE1* is a key regulator of the ECM that has been associated with recurrent pregnancy loss.^25^ This prompted us to investigate whether other genes associated with pregnancy loss or stillbirth, a common outcome of perinatal syphilis,^1^ were also dysregulated by *T. pallidum*. Indeed, assembled heatmaps of expression of genes either causally associated with stillbirth or those that have been identified as candidates for involvement in stillbirth (**Fig. S1**).^26^ While many of the genes were differentially regulated between the three cell lines, some, including *NOL6, NUP98,* and *PRMT5*, exhibited a similar pattern of *T. pallidum*-mediated downregulation. Although these findings do not imply causality, loss-of-function variants in these genes have been reported in stillbirth cases,^26^ raising the possibility that their reduced expression during infection may represent a mechanistic link between *T. pallidum*, placental dysfunction and fetal loss that warrants further investigation.

## Discussion

Despite its susceptibility to penicillin, syphilis has continued to cause substantial disease in humans for hundreds of years.^27^ A significant hurdle in syphilis research has been historical difficulty culturing *T. pallidum in vitro*. Discovery of culture conditions that permit *T. pallidum* replication in cottontail rabbit Sf1Ep cells in 2018 was a significant advancement for the field;^11^ however, a paucity of similar human-based models has prevented mechanistic studies that could advance our understanding of *T. pallidum* pathogenesis in human cells. Here, we identify human placental trophoblast cell lines that support sustained *T. pallidum* replication at rates comparable to Sf1Ep cells. These models provide the first human trophoblast cellular systems for long-term propagation of *T. pallidum* and establish a tractable platform to interrogate host-pathogen interactions at the maternal-fetal interface.

We found that three trophoblast cell lines, JEG-3, BeWo, and HTR-8, facilitated successful *T. pallidum* replication. All three of these cell lines have been widely used to investigate pathogenesis of other pathogens that are able to infect and cross the placenta.^17–21^ However, due to the historical inability to culture *T. pallidum* continuously in human cell lines, studies using these cell lines to study congenital syphilis are severely limited. While two prior studies used BeWo cells to investigate syphilis in the placenta, they used either recombinant expression of *T. pallidum* outer membrane proteins either alone^28^ or in the Lyme disease-causing spirochete *Borrelia burgdorferi,*^29^ and did not examine *T. pallidum* replication. As demonstrated by our findings that transcriptional changes were absent from trophoblasts co-cultured with heat-inactivated *T. pallidum*, spirochete replication and metabolic activity has significant implications for host trophoblast cells, highlighting the importance of examining infection-mediated changes in a replication-competent model.

A recent preprint seeking to identify human cell lines that support *T. pallidum* assessed replication in two placental cell lines, JAR and JEG-3, which they reported to be “non-supportive” of replication due to low bacterial counts.^30^ They also did not examine any downstream implications for trophoblast function or gene expression The discrepancy between these results and our successful sustained co-culture of JEG-3 with *T. pallidum* could lie in strain-specific differences, as we evaluated growth of the Nichols strain while Capuccini *et al* assessed the SS14 strain. However, we have found that Mexico A, a *T. pallidum* strain from the SS14-like cluster,^31^ is well supported in JEG-3 cells, and that BeWo cells also support SS14 replication, in ongoing studies. By defining conditions that enable efficient, long-term spirochete replication in three trophoblast cell lines, our study establishes a foundation for mechanistic dissection of *T. pallidum* interactions with placental cells.

The ability of *T. pallidum* to replicate efficiently in all three trophoblast lines examined supports the concept that placental cells provide a permissive niche for this highly host-dependent spirochete. Interestingly, *T. pallidum* replication rates differed slightly among the three cell lines, with fastest doubling times occurring in HTR-8 cells. The reason for these differences is unclear, but one possibility is that EVT-like trophoblasts possess features that are particularly permissive for *T. pallidum* growth. Indeed, other placental TORCH pathogens including the bacterium *Listeria monocytogenes*^32,33^ and parasite *Toxoplasma gondii*^34^ show preferential invasion and replication in EVTs versus other trophoblast cell types. Further studies using additional EVT models will be needed to determine whether EVT biology and immune-privileged status enhances *T. pallidum* replication.

We also observed that faster *T. pallidum* doubling times correlated with increased numbers of transcriptional changes, with BeWo and HTR-8 having both lower spirochete doubling times and higher numbers of significantly dysregulated transcripts than JEG-3. Together with the absence of comparable transcriptional changes in cultures exposed to heat-inactivated *T. pallidum*, these findings suggest that the trophoblast transcriptional response is linked to active spirochete growth, metabolism, and replication rather than passive exposure to bacterial material. This interpretation is consistent with the highly reduced metabolic capacity of *T. pallidum* and its reliance on host-derived nutrients for survival,^35^ which likely contributes to transcriptional changes in permissive host cells. This also underscores the importance of examining the impact of *T. pallidum* infection on trophoblast cells in replication-competent models, as recombinant or heterologous expression systems may not fully recapitulate *T. pallidum*-mediated effects on host cells.

Using our replication-competent trophoblast models, we identified several pathways that were disrupted by *T. pallidum* across all three placental cell lines. The first was cholesterol metabolism, with trophoblasts demonstrating transcriptional changes consistent with increased synthesis/uptake and reduced efflux of cholesterol. This is highly noteworthy given that *T. pallidum* has been shown to use cholesterol-dependent lipid raft-mediated endocytosis to cross endothelial barriers.^36^ Thus, if a similar mechanism occurs to bypass the placental barrier, increased production and retention of intracellular cholesterol could potentially facilitate spirochete invasion. Conversely, host environment-derived cholesterol is a major constituent of the lipids found within *T. pallidum*.^37,38^ Cholesterol retention within host cells could therefore function as a protective mechanism by limiting bacterial access to this resource, whereas increased cholesterol biosynthesis or trafficking may create a metabolic environment that favors replication. Future studies that perturb cholesterol synthesis, storage and transport will be needed to determine whether these responses restrict or support *T. pallidum* growth in placental cells.

In addition to changes in cholesterol metabolism, we observed transcriptional changes consistent with deficient type I IFN signaling and significant downregulation of ISGs across all trophoblast cell lines. Type I IFN signaling and subsequent upregulation of ISGs plays an important antiviral role in the placenta, with deficient signaling facilitating increased transplacental transmission of viral TORCH pathogens, such as Zika virus.^39^ While generally considered protective against viral infection, type I IFN signaling has conflicting and pathogen-specific roles during bacterial infection;^40^ whether activation of type I IFN signaling is protective or detrimental during *T. pallidum* infection is unclear. A previous study reported increased ISGs in peripheral blood mononuclear cells (PBMCs) from patients with secondary syphilis,^41^ including increases in *OAS3* and *MX1*, both of which were decreased in our study. In contrast, endothelial cells exposed to *T. pallidum in vitro* for up to 72 hours exhibited reduced expression of interferon regulatory factor 1 (IRF1) and downstream ISGs,^42^ suggesting that the impact of *T. pallidum* on type I IFN signaling may be both tissue and time-dependent.^35^ While it is unknown whether ISGs protect against or facilitate *T. pallidum* invasion and replication in the placenta, reduced type I interferon signaling may create an immunologically permissive state that facilitates spirochete persistence and replication in trophoblasts. Moreover, because regulation of interferon activity is important for placental development, immune defense, and pregnancy maintenance, attenuation of these pathways could have consequences not only for bacterial persistence but also for placental function and pregnancy outcome.^43^

Finally, pathway enrichment analysis suggested dysregulation of extracellular matrix (ECM) organization in trophoblasts cultured with *T. pallidum*. This is consistent with transcriptional responses reported in brain endothelial cells^42^ and prior evidence that *T. pallidum* can bind to^44–46^ and even degrade^46^ various components of the ECM. In particular, we observed significantly increased expression of fibronectin, a known binding target of *T. pallidum*.^45^ While binding of *T. pallidum* to fibronectin has been demonstrated in various human cell types, such an interaction has not been investigated in the placenta. It would therefore be interesting to determine whether the increased fibronectin in infected trophoblasts plays a role in spirochete adherence.

Notably, we also uncovered transcriptional changes in genes either causally associated with stillbirth or proposed as candidate contributors to stillbirth. The strongest evidence for this association was significantly increased expression of *SERPINE1*, a key regulator of ECM remodeling whose increased expression is associated with recurrent pregnancy loss.^25^ Several other genes, including *NOL6*, *NUP98* and *PRMT5*, exhibited conserved trends towards *T. pallidum*-associated downregulation across trophoblast cell lines. Because loss-of-function variants in these genes have been reported in stillbirth cases,^26^ their reduced expression during trophoblast co-culture may represent a potential link between *T. pallidum* infection, altered placental cell function, and adverse pregnancy outcomes. Future studies will be needed to determine whether these transcriptional changes contribute directly to placental dysfunction or fetal loss during congenital syphilis.

In sum, we identify human placental trophoblasts as permissive host cells that support efficient, sustained *T. pallidum* replication *in vitro*, establishing the first replication-competent human models of *T. pallidum* infection at the maternal-fetal interface. By integrating long-term co-culture with transcriptomic profiling, we uncover conserved trophoblast responses involving cholesterol metabolism, extracellular matrix organization, and attenuated interferon signaling. These pathways provide mechanistic entry points to investigate how *T. pallidum* adheres to, remodels, and persists within placental cell environments. More broadly, these models create a tractable platform for defining the cellular basis of congenital syphilis, identifying biomarkers of placental infection, and testing interventions aimed at limiting spirochete replication and transplacental transmission. These systems therefore provide a timely experimental foundation for addressing the biological mechanisms underlying the escalating congenital syphilis epidemic.

## Methods

### Cell Culture

The human trophoblast cell lines JEG-3 (ATCC HTB-36), BeWo (ATCC CCL-98) and HTR-8/SVNeo (ATCC CRL-3271) were obtained from ATCC and maintained in DMEM/F-12 (Gibco, 11330032) supplemented with 10% fetal bovine serum (FBS) (Sigma, F135). The cottontail rabbit cell line Sf1Ep (ATCC CCL-68) was obtained from ATCC and maintained in Eagle’s MEM with nonessential amino acids, L-glutamine, sodium pyruvate, and 10% heat-inactivated FBS.^11^ All cell lines were cultured at 37°C and 5% CO_2_ prior to inoculation with *T. pallidum*.

### In vitro Cultivation of T. pallidum in Sf1Ep

*T. pallidum* subspecies *pallidum* Nichols was obtained from J.N. Miller at the UCLA Geffen School of Medicine. This strain was originally isolated from the cerebrospinal fluid of a neurosyphilis patient in 1912.^47^ A recently generated GFP-tagged *T. pallidum* Nichols strain^22^ was obtained from M.J. Caimano at University of Connecticut. Both strains were maintained in Sf1Ep cells *in vitro* as previously described.^11,48^ Briefly, *T. pallidum* was cultured with Sf1Ep cells in TpCM2 medium at 34°C in a low-oxygen incubator with 1.5% O_2_, 5% CO_2_, and 93.5% N_2_. Cultures were passaged once a week by adding inoculum containing 2-5 million spirochetes to newly seeded Sf1Ep in fresh medium.

### Cultivation of T. pallidum in Placental Cell Lines

JEG-3, BeWo, and HTR-8 cells were maintained in standard culture conditions prior to inoculation with *T. pallidum.* One day prior to inoculation, each cell line was seeded in 6-well plates at a density of 50,000 cells per well. TpCM2 medium was prepared as described^48^ and transferred to the low-oxygen incubator for equilibration overnight. On the day of inoculation, existing culture media from seeded cells was removed. Cells were washed with 1 mL pre-equilibrated TpCM2, replenished with 4 mL fresh TpCM2, and moved to the low oxygen incubator for 3-4 hours of equilibration.

To prepare inoculum, conditioned TpCM2 medium from Sf1Ep cultures containing *T. pallidum* was collected. Sf1Ep cells were trypsinized with trypsin-EDTA (Millipore Sigma, T4049) for 5 minutes at 37°C and combined with collected culture medium. Suspensions were centrifuged at 125 *g* for 5 minutes to pellet Sf1Ep cells. To eliminate any remaining Sf1Ep cells, supernatants were gravity filtered through a 0.5 µM syringe filter with an SFCA membrane (Corning, 431220) to generate inoculum containing pure *T. pallidum*. After counting with darkfield microscopy, inoculum containing 2-5 million spirochetes was added to each well of 6-well plates seeded with JEG-3, BeWo, or HTR-8 cells equilibrated in TpCM2. Inoculated cultures were maintained at 34°C in a low-oxygen incubator with 1.5% O_2_, 5% CO_2_, and 93.5% N_2_ for 6-7 days, at which point they were passaged as described above for Sf1Ep.^48^

### Confocal microscopy

For imaging experiments, 0.5×10^5^ cells (Sf1Ep, HTR-8, BeWo, or JEG-3) were seeded overnight on sterile 25 mm diameter poly-L-lysine-coated 1.5 glass coverslips (Neuvitro, GG251.5PLL), as described previously, and inoculated with GFP-expressing *T. pallidum* (3.4×10^6^ cells + 200 µg/ml kanamycin). After 7 days of infection, cells were washed once with PBS and fixed with 3.7% paraformaldehyde for 10 minutes at room temperature. After three additional PBS washes, fixed cells were permeabilized for 10 minutes with PBS containing 0.1% Triton X-100, followed by a wash with fresh PBS. DNA and F-actin were stained during a 10-minute incubation in PBS containing Hoechst 33342 (1:10,000; Invitrogen, H1399) and AlexaFluor™ 568 phalloidin (1:500; Invitrogen, A12380), respectively. Cells were washed three more times with PBS before mounting the coverslip onto glass slides with ProLong Gold Antifade Mounting Medium (Invitrogen, P36934). The following day, the coverslips were sealed with clear nail polish and imaged using the Olympus Fluoview FV3000 confocal microscope equipped with the Fluoview FV315-SW software. Using a step size of 0.20 µm, z-stacked images were acquired and processed with Olympus cellSens Dimension software.

### RNAseq

Total RNA was collected from cells inoculated with *T. pallidum* for 3 or 7 days and corresponding controls using an RNeasy kit (Qiagen, 74182) followed by on-column DNase treatment with an RNase-free DNase kit (Qiagen, 79254). Poly-A RNA sequencing was conducted by the Genomic and RNA Profiling Core at Baylor College of Medicine. Libraries were prepared using a TruSeq Stranded mRNA Kit (Illumina, 20020594/5) and sequenced on an Illumina NovaSeq X instrument with a 10B flow cell. Sequencing was carried out in paired-end 150 bp mode, generating approximately 30-40 million reads per sample. Raw FastQ data reads were subjected to quality control with FastQC, trimmed with TrimGalore v0.6.7, and mapped to the human hg38 reference genome (Ensembl GRCh38.101) using STAR v2.7.10b. Protein-coding transcripts were extracted from the raw counts matrices for each biological triplicate, and transcripts with zero counts across all samples were removed. Differential expression analyses comparing control and *T. pallidum*-infected cells within each cell line at either 3 or 7 days were performed using PyDESeq2 v0.5.2. Initial filtering for baseMean >= 10 removed transcripts with extremely low expression. Significantly upregulated or downregulated transcripts were defined as p<0.05 and log_2_FoldChange > 0.5 or < -0.5 (approximately > 1.4-fold change). Differentially expressed genes were used for principal component analysis (PCA) in Scanpy v1.11.4, and PCA and volcano plots were generated using Seaborn v0.13.2. Genes that were significantly dysregulated across all three cell lines at a particular timepoint were subjected to pathway enrichment analysis using Gene Ontology Biological Processes via Enrichr.^49–51^

### qRT-PCR Validation

RNA used for validation was the same RNA analyzed by RNAseq, with an additional group of cells co-cultured with heat-inactivated (dead) *T. pallidum* as another control. Either 500 ng (JEG-3 and HTR-8) or 200 ng (BeWo) of DNAse-treated total RNA was reverse-transcribed using SuperScript II Reverse Transcriptase (Invitrogen, 18064-014). cDNA was amplified and quantified using SsoAdvanced Universal SYBR Green Supermix (BioRad, 1725274) with primers listed in **Supplementary Table 1**. Relative mRNA expression was determined using the comparative threshold cycle method (ΔΔC_t_) method and normalized to expression of *YWHAZ* (JEG-3 and HTR-8) or *TOP1* (BeWo).

### Statistical Analysis

All experiments were conducted in three biological replicates. For qRT-PCR, statistical analysis was conducted in GraphPad Prism 10.6.1. Data is shown as mean ± standard error of the mean. Significance (p<0.05) was assessed using one-way ANOVAs with Tukey’s multiple comparisons.

## Supporting information

Supplemental materia

## Acknowledgements

This project was supported by an NIH/NIAID R21AI194017 to IUM and DGE, NIH/NIAID R01AI176505 to IUM, and 2R01AI141958 to DGE. ERM was supported by a Postdoctoral Fellowship in Infection and Immunity from Baylor College of Medicine. This project was also partly supported by the Genomic and RNA Profiling Core at Baylor College of Medicine with funding from the NIH S10 grant (1S10OD036427). Figure 5M was generated using Biorender.com. GFP-expressing *T. pallidum* were a generous gift from Dr. Melissa Caimano (UConn Health).

## Disclosures

The authors have no conflicts of interest to disclose.

## Notes

### Competing Interest Statement

The authors have declared no competing interest.

