## Supplemental materia for "Human placental trophoblasts support sustained *Treponema pallidum* replication and reveal new candidate host pathways implicated in congenital syphilis": Supplementary Figure 1.pdf

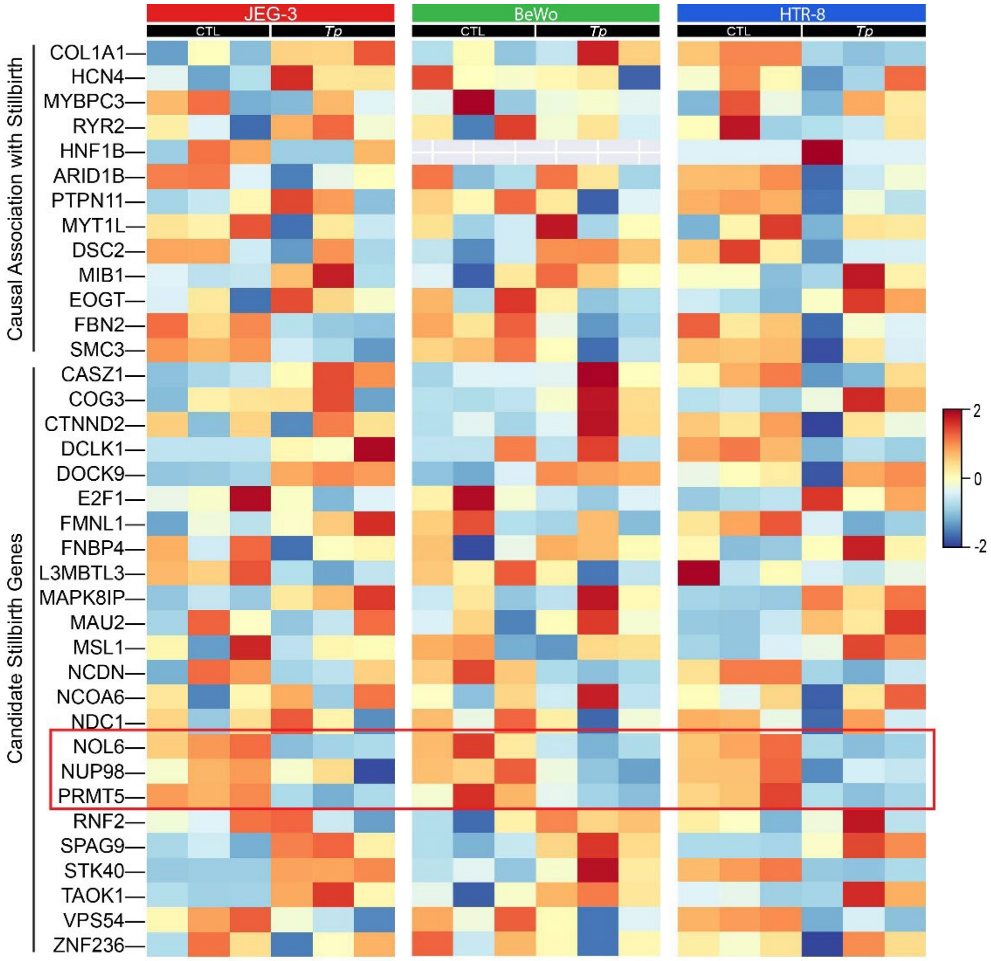


**Supplementary Figure 1: Heatmap of transcript levels of genes associated with stillbirth.** Transcript levels were determined by RNAseq in each cell line after 7 days of co-culture with *T. pallidum* (*Tp*). Legend refers to Z-score, with red indicating increased transcript levels and blue indicating decreased transcript levels. The red box highlights three genes, *NOL6, NUP98,* and *PRMT5*, which show a conserved trend of reduced transcript levels in *T. pallidum*-infected cells.
