## Supplemental materia for "Human placental trophoblasts support sustained *Treponema pallidum* replication and reveal new candidate host pathways implicated in congenital syphilis": Supplementary Table 1.pdf

**Supplementary Table 1:** Primers used for qRT-PCR. Sequences are reported 5' to 3'.

| Gene | RefSeq | Forward | Reverse |
|---|---|---|---|
| *ABCA1* | NM_005502.4 | AGTATGGAAGAATGTGAAGC | ATAACCATCTCCAAACCTAT |
| *ADAP2* | NM_018404.3 | GGATCTTCATCTGTCTCAAC | GTGGATCATAAACTCCACA |
| *FN1* | NM_212482.4 | AATCCAAATGCCTCTACA | AGGAACCCTGAACTGTAA |
| *HMGR* | NM_000859.3 | GATCTGGAGGATCCAA | ATTCATAATTCCAACCACA |
| *IFITM2* | NM_006435.3 | GGAAGTGGCTATGCTG | TCCTGTCCCTAGACTTCA |
| *IFITM3* | NM_021034.3 | CTCCGTGAAGTCTAGGG | GCCTCCTGATCTATCCA |
| *LDLR* | NM_000527.5 | GCTACCCCTCGAGACA | CTTTGGTCTTCTCTGTCTTT |
| *MX1* | NM_002462.5 | ACCATTCCAAGGAGGT | AGCTCTAATATACGTTCCTTC |
| *OAS3* | NM_006187.4 | TTGATACCATCTGTTCATTT | AGAGCCACCCTTGAC |
| *SERPINE1* | NM_000602.5 | CTCTGAGAACTTCAGGATG | CATAGGGTGAGAAAACCAC |
| *TOP1* | NM_003286.4 | GATGAACCTGAAGATGATGGC | TCAGCATCATCCTCATCTGC |
| *YWHAZ* | NM_001135699.2 | TGTAGGAGCCCGTAGGTCATC | GTGAAGCATTGGGGATCAAGA |
